# An exploratory study on the role of the stiffness of breast cancer cells in their detachment from spheroids and migration in 3D collagen matrices

**DOI:** 10.1101/2021.01.21.427639

**Authors:** Ghodeejah Higgins, Jessica E. Kim, Jacopo Ferruzzi, Tamer Abdalrahman, Thomas Franz, Muhammad H. Zaman

## Abstract

**Background:** Tumour-cell detachment is a critical early event in the metastatic cascade. However, the role of the cell’s mechanical properties in detachment and migration is not well understood. This exploratory study aimed to assess how intracellular stiffness changes these processes.

**Methods:** MDA-MB-231 cells were embedded as 10,000-cell spheroids in 2 and 4 mg/ml collagen matrices. Intracellular stiffness was assessed with mitochondria tracking microrheology of cells that migrated distances equivalent to four and six times the cell diameter (d_C_) from the spheroid and compared to cells at the spheroid surface (0d_C_), representing medium, high and no migration, respectively.

**Results and discussion:** The mitochondrial mean square displacement and intracellular stiffness decreased during detachment and migration for both collagen concentrations (i.e. rigidities). The mean square displacement of 4d_C_ and 6d_C_ cells was similar, whereas cell stiffness was lower for 4d_C_ than for 6d_C_ cells. With increasing matrix rigidity, the intracellular stiffness decreased for 0d_C_ cells and did not change for 4d_C_ and 6d_C_ cells. It is proposed that decreased cell stiffness drives detachment and migration and increased matrix rigidity physically hinders migration, and cells need to become softer or remodel the environment to migrate. The independence of the stiffness of migrated cells from matrix rigidity suggests that cells remodel their environment through matrix proteins cleavage to migrate.

**Conclusions:** The study revealed the collective effects of enhanced migratory conditions and increased matrix rigidity on the mechanical properties of the cells. The expression of matrix metalloproteinases and transforming growth factor β and the role of cell volume on detachment and migration in matrices with varying pore sizes are proposed targets for further studies on metastatic cancer cells.

## 1. Introduction

Metastatic cancer cells alternate between various labour-intensive tasks associated with disease progression, including detachment from the tumour, invasion into the tumour environment and formation of new tumours [1]. Considerable attention has focussed on understanding the role that the mechanical properties of cancer cells play in these processes. It has been shown that the mechanical properties have crucial implications in regulating chemical and mechanical cues and in maintaining cell or tissue architecture in several cell types, including metastatic cancerous cells [2, 3].

Many of these studies have probed cells in two-dimensional environments. However, cells in two-dimensional (2D) and three-dimensional (3D) environments exhibit substantial differences in morphology, gene expression, and mechanical and structural properties. Several studies reported a marked decrease in intracellular fluctuations from 2D to 3D environments [3, 4]. Additionally, 3D *in vitro* systems can recapitulate *in vivo* conditions more closely than 2D systems [4].

Therefore, studies of solid tumours should incorporate 3D models that mimic complex tumour architecture, including cell-cell and cell-matrix interactions [5]. Cell spheroids are powerful in mimicking many aspects of solid tumours, including architecture (i.e. an outer proliferative layer of cells, a middle layer of senescent cells and a core of necrotic cells), enhanced cell-cell and cell-matrix connections, and gene expression [6, 7].

Tumour spheroid assays have addressed essential questions relating to spheroid morphology, growth, viability, drug development [5], signalling pathways associated with cell detachment and invasion [8], cancer stem cell content [9]. However, few studies have investigated cell mechanics in the detachment and invasion process due to the technical challenges of quantifying the mechanics of live cells in 3D environments during detachment and invasion. Microrheology can offer non-invasive techniques that allow passively probing intra-cellular mechanics to characterise single and collective cell behaviour without introducing cellular and matrix deformation [3].

It is generally accepted that high tissue rigidity is a characteristic of solid tumours [10] and that cells adapt their stiffness to environmental stiffness [11]. It is essential to investigate whether and how cells adapt their stiffness during tumour detachment and invasion as the disease progresses. It has been shown that reduced cell-cell adhesion decreases cell stiffness [12] and increases migration [13] and that increased migration and decreased cell stiffness are directly correlated [14, 15]. However, it is not yet known whether these observations of cellular mechanics translate directly to physiologically relevant environments that mimic the tumour-cell detachment process. Considering that the mechanical properties of cells depend on the properties of the extracellular matrix (ECM), it is essential to clarify how cells regulate their mechanical properties when the properties of the ECM change.

This study aimed to explore the role of intracellular stiffness of metastatic breast cancer cells during detachment from spheroids and invasion, and to identify targets for further research to complement this initial work. Different levels of migratory capacity were established by considering cells increasingly migratory with increasing migration distance from the spheroid.

## 2. Materials and methods

### 2.1. Cell culture

Metastatic breast adenocarcinoma cells MDA-MB-231 (purchased from the American Type Culture Collection, ATCC, Manassas, VA) were cultured in DMEM (Dulbecco’s Modified Eagle Medium, Life Technologies), supplemented with 10% foetal bovine serum (FBS) and 1% penicillin-streptomycin, and maintained in 25 cm^2^ tissue flasks (Costar, Corning Life Science, Acton, MA) at 37°C and 5% CO_2_ until near-confluency. Media was replaced every 3-4 days. Cells were used in passages 6-16.

### 2.2. Spheroid formation

Cell suspensions were grown in agarose-coated well plates to form spheroids according to the liquid overlay method [16]. An agarose solution was prepared by diluting 0.18 g agarose (Sigma-Aldrich, St. Louis, MO) in 12 ml 1XPBS and heated in a microwave oven for 36 s or until the agarose dissolved. The solution was immediately transferred to sterile conditions and kept on a hot plate to prevent premature gelation. A 96-well plate was prepared by pipetting 70 μl of the hot liquid agarose solution onto the flat bottoms of the wells, ensuring the solution covered the entire surface and contained no air bubbles. The agarose-coated 96-well plate was exposed to ultraviolet light for 30 minutes for sterilisation. Separately, cell monolayers were lifted from the culture flask using standard laboratory trypsinisation procedures. Spheroids of 10,000 cells were created by placing 100 μl of trypsinised cells (100,000 cells/ml containing 2.5% Matrigel) into the agarose-coated wells, centrifuged at 1,000 rpm for 5 mins, and incubated at 37°C and 5% CO_2_ for 72 hrs.

### 2.3. Spheroid collagen embedding

MDA-MB-231 spheroids were embedded in Type I Rat Tail Collagen Solutions (BD Biosciences) at 2 and 4 mg/ml final collagen concentrations. Type I Rat Tail Collagen at a stock solution of 9.61 mg/ml was combined with equal amounts of neutralising solution (100 mM HEPES in 2X PBS with pH 7.3) and further diluted with 1XPBS on ice to create 2 and 4 mg/ml collagen solutions. Spheroids were added to the unpolymerised collagen solutions on 35mm glass-bottom dishes, with each 50 μl collagen solution containing one spheroid to avoid spheroid-spheroid interaction, and incubated at 37°C and 5% CO_2_ for 1 hr. After 1 hr, the collagen solutions had polymerised, and 500 μl media were added.

### 2.4. Mitochondrial particle tracking microrheology

The use of mitochondria as tracer particles to determine the viscoelastic response of cells has been validated by Mak, Kamm (3), who showed that the mean square displacement (MSD) of mitochondria and ballistically injected nanoparticles was similar when used in metastatic breast cancer MDA-MB-231 cells.

Embedded MDA-MB-231 tumour spheroids were prepared for mitochondrial tracking by adding 500 nM Mitotracker Red solution (Life Technologies, Carlsbad, CA) to the supplemented DMEM media 24 hrs before experimentation to allow the Mitotracker solution to sufficiently diffuse through the spheroid. Before imaging, the media was removed from the wells, and the wells were placed in an environmental chamber at 37°C and 5% CO_2_.

After 24 hrs, cells started detaching from the spheroid and migrating into the collagen gel predominantly as linear branches with a leader cell and follower cells.

To avoid boundary effects of the rigid 2D glass-bottom surface and to ensure full engulfment, cell branches near the middle of the collagen gel were located and imaged. Mitochondrial fluctuations of branching cells at the spheroid surface and varying distances from the spheroid were captured using time-lapse fluorescent imaging for 120 s (exposure time: 50 ms/frame; spinning disk confocal microscope, 63x 1.4 oil immersion objective and CCD camera, Hamamatsu Photonics, Hamamatsu, Japan). Fluorescent images were acquired of surface and migrated cells (branched). The distance migrated (d_m_) from the spheroid was reported in cell diameter (d_C_), where d_m_ = 0d_C_, 4d_C_ and 6d_C_ correspond to the cell at the spheroid surface, intermediately migrated, and furthest migrated, respectively. These experiments were repeated on two independent days (n = 2), with data taken from at least two spheroids, each with at least five branches, resulting in 10 cells per condition. Each branch comprised at least six migrated cells. Cells with at least 60 mitochondria, visible for at least 80% of the total image acquisition time, were retained for analysis.

Post-processing involved constructing mitochondrial trajectories using TrackMate [17] in Fiji Image J [18]. The tracking assumed particles with a diameter of 1 μm and allowed up to 4 missed frames between subsequent spots to account for missing detections. Mitochondrial trajectories were imported into MATLAB (The MathWorks, Natick, MA), and the ensemble-averaged, time-dependent MSD was calculated using @msdanalyzer [19] according to:

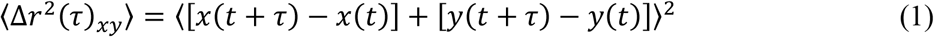

Here, 〈Δ*r*^2^ (*τ*)_*xy*_〉 is the ensemble-averaged MSD, τ is the time interval or delay time between the first and last image frame used for the analysis, and *x(t)* and *y(t)* refer to the spatial coordinates of particle positions at time *t*.

For viscoelastic materials, the MSD scales nonlinearly with the delay time τ according to a power-law relationship, 〈Δr^2^ (τ)〉 ~ τ^α^ [20]. The power-law coefficient α = ∂ln 〈Δr^2^ (τ)〉/∂ln (τ) represents the slope of the logarithmic MSD-τ curve.

### 2.5. Statistical analysis

Data are reported as mean ± standard deviation in the text. Error bars in graphs represent the standard error of the mean (SEM). Two-way ANOVA was employed to detect differences in MSD and α of cells at different delay times. ANOVA assessed two main effects, i.e. distance from the spheroid and changes in collagen concentration, and one interaction between main effects. The migrated distance contained three levels, i.e. no migration, medium migration, and large migration, with which posthoc analysis was performed using Tukey’s HSD test. The collagen concentration contained two levels, i.e. 2 mg/ml and 4 mg/ml collagen, and ANOVA was followed up with linearly independent pairwise comparisons among estimated marginal means.

A criterion for statistical significance of *p* < .05 formed the basis of all evaluations. SPSS Statistics for Windows (Version 25.0, IBM Corp., Armonk, NY) was used for all statistical analyses.

## 3. Results

To provide a platform for cell detachment from tumours and migration in physiologically relevant environments, metastatic breast cancer cells (MDA-MB-231) were embedded as tumour spheroids in 3D collagen matrices. Mitochondrial particle tracking microrheology experiments (n = 2) were conducted after 24 hrs of incubation when cells started detaching from spheroids.

A low MSD value corresponds to constrained particle fluctuations and indicates a stiffer, solid material, whereas a high MSD indicates greater particle motility and a more fluid-like material. For short delay times, the mitochondrial MSD primarily represents the viscoelastic intracellular properties, whereas active intracellular motor-driven processes dominate the MSD for long delay times [3, 21]. The power-law coefficient α helps classify the motion of the tracer particles. An α close to 1 corresponds to diffusive motion such as thermal fluctuations in Newtonian fluids, whereas an α close to 0 indicates constrained, sub-diffusive motion such as thermal fluctuations in an elastic material [21, 22].

Following detachment from the tumour surface, cells invaded the tumour environment predominantly in linear branches. Several cells trailed a leader cell in these branches and migrated away from the tumour (Figure 1).

**Figure 1:**
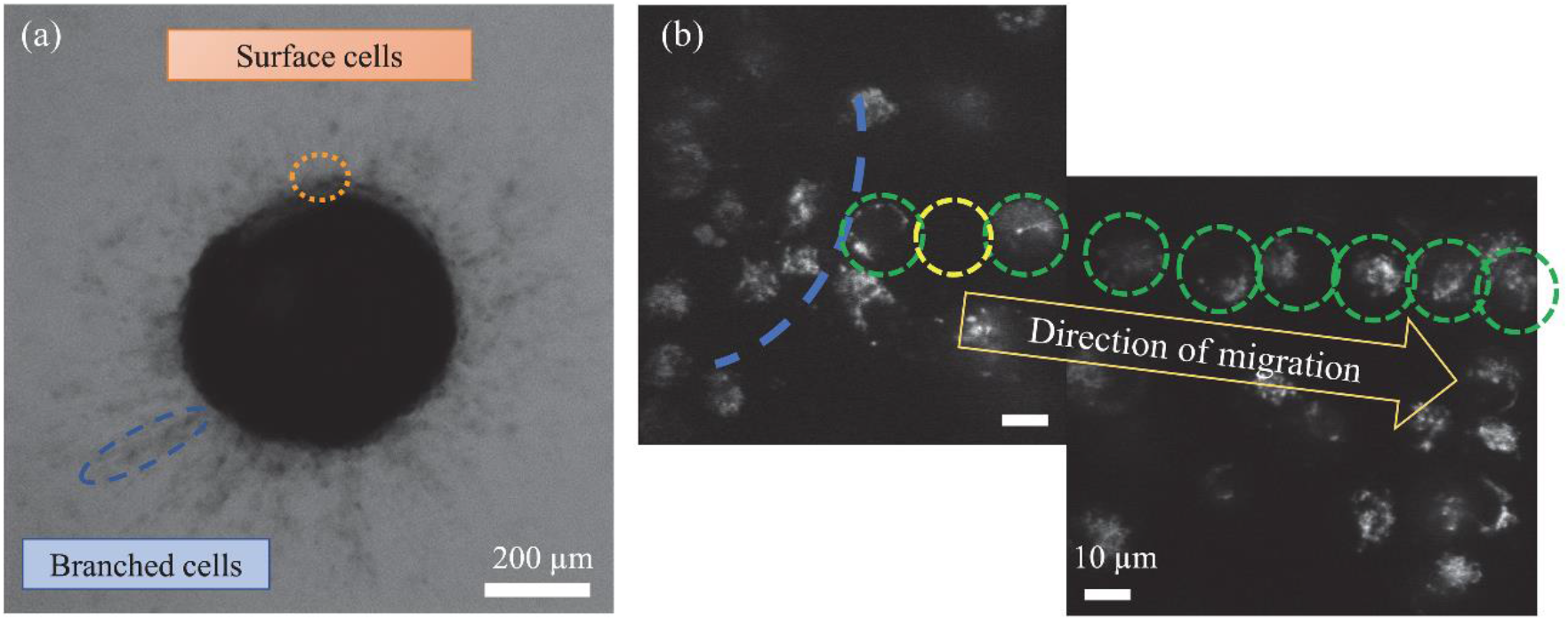
Illustration of detaching (surface) and migrated (branched) cells as (a) bright-field image (5x magnification) and (b) mitochondrial fluorescent image (63x magnification). The blue dashed line indicates the spheroid border, green dashed circles indicate the approximate diameter of cells migrating away from the spheroid in a branch, and the yellow dashed circle indicates a cell out of focus.

### 3.1. Effect of detachment and migration: Migrated cells are more deformable than detaching cells

The effect of detachment and migration of cells on mitochondrial fluctuations and cell stiffness was assessed in 2 and 4 mg/ml collagen.

The MSD of 0d_C_, 4d_C_, and 6d_C_ cells in 2 and 4 mg/ml collagen increased with increasing delay time (Figure 2). Lower and flat MSD-τ curves indicate solid-like intracellular behaviour, whereas higher and increasingly sloped curves indicate fluid-like, intracellular motor-driven mechanics. MSD and α were further compared for short delay times of τ = 0.05 s and 2 s, respectively, to determine the viscoelastic properties of cells as the distance from the spheroid increased (Figure 3).

**Figure 2:**
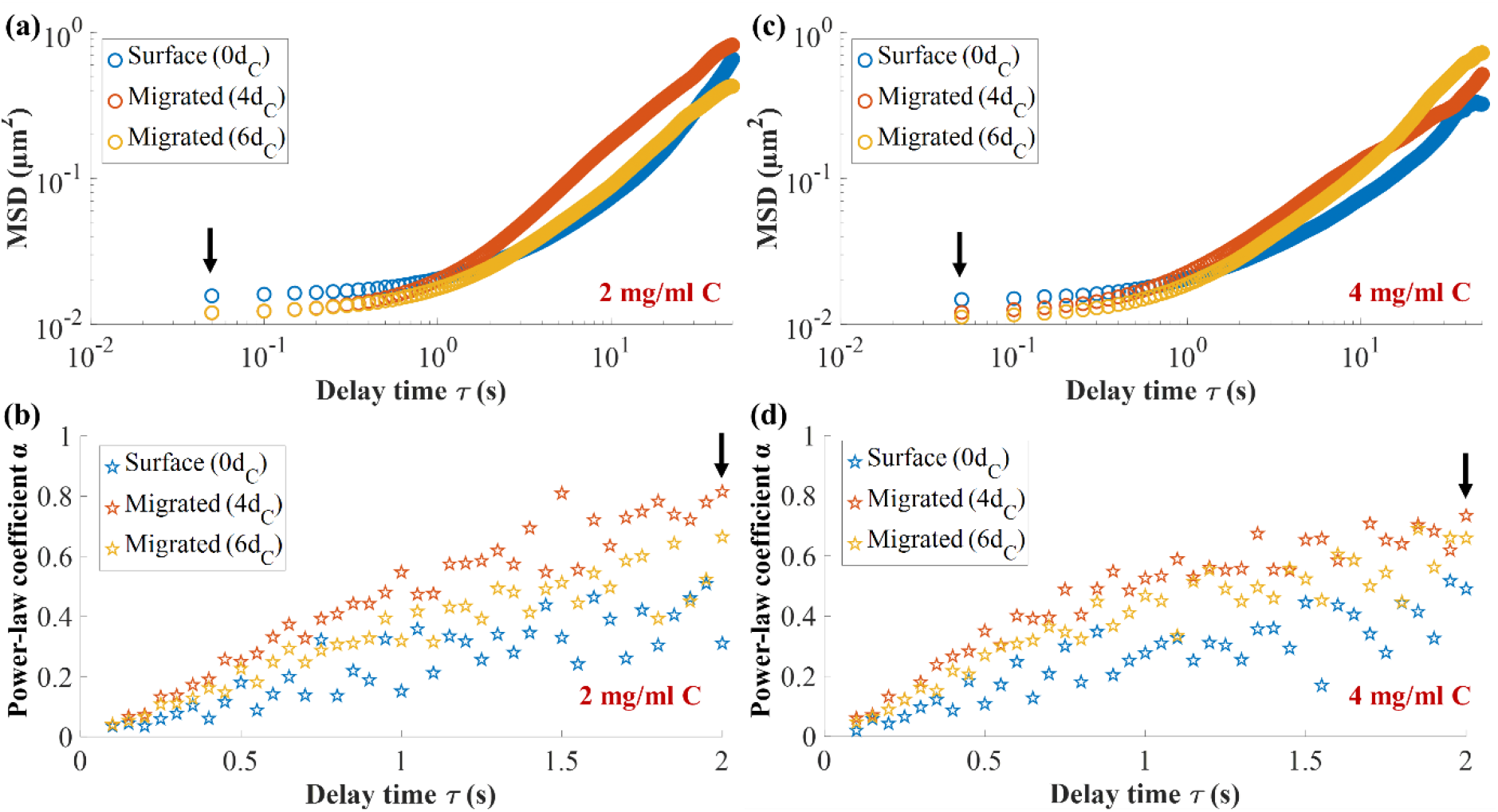
Intracellular fluctuations from spheroids (n = 2) in 2 and 4 mg/ml collagen gels. MSD and power-law coefficient a versus delay time of cells in 2 mg/ml collagen (a, b) and 4 mg/ml collagen (c, d). Error bars omitted for clarity. Arrows indicate the delay time of τ = 0.05 s for MSD and τ = 2 s for α. Trends of MSD and α of cells were similar in 2 and 4 mg/ml collagen. Mitochondrial fluctuations of 0d_C_ were larger than migrated 4d_C_ and 6d_C_ cells. 0d_C_ cells exhibited the largest cell stiffness, followed by 6d_C_ and then 4d_C_ cells.

**Figure 3:**
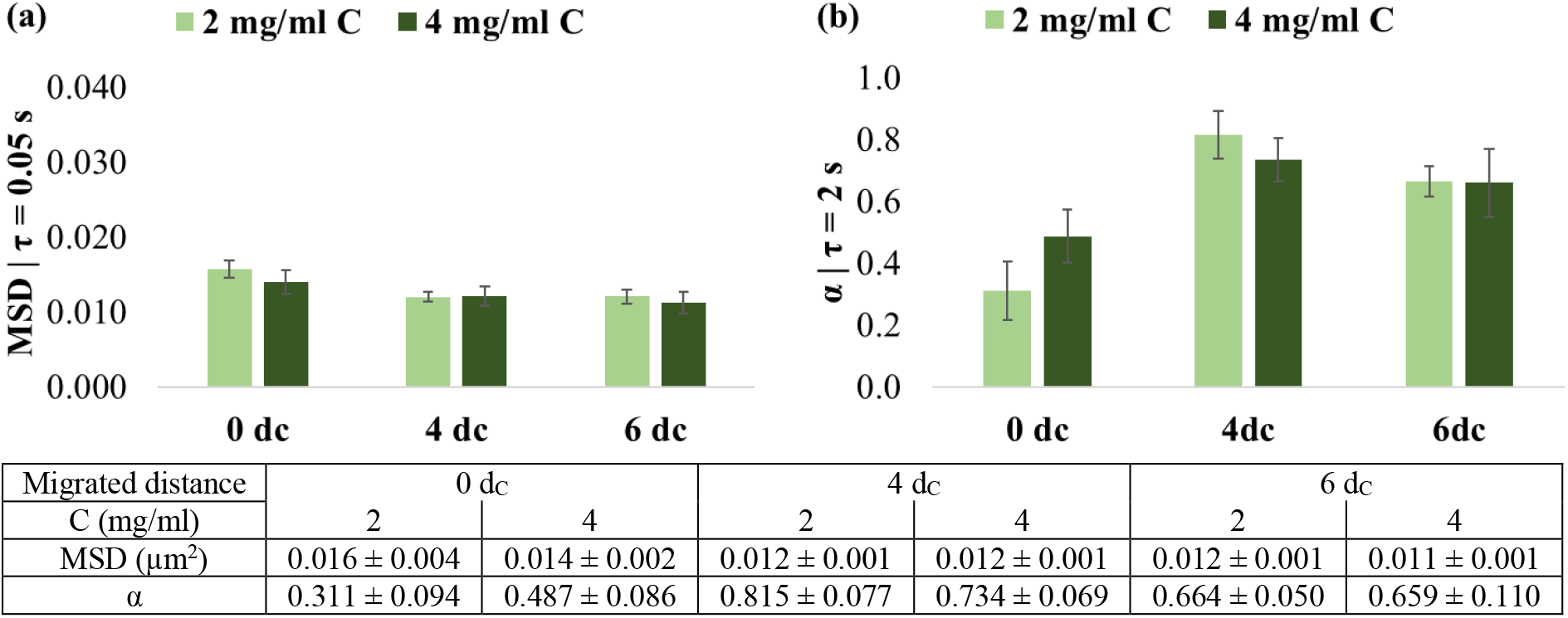
Comparison of (a) mean square displacement (MSD) and (b) power-law coefficient a of cells at the spheroid surface (0d_C_) and after migration of a distance of 4d_C_ and 6d_C_ from the spheroid surface in 2 and 4 mg/ml collagen matrices (C) (d_C_ indicates cell diameter). All differences were nonsignificant. Values are mean ± SEM; error bars are SEM.

In 2 mg/ml collagen matrices, mitochondrial fluctuations (at τ = 0.05 s) were larger and the power-law coefficient α (at τ = 2 s) was smaller for cells at the spheroid surface, i.e. d_m_ = 0d_C_, than for cells that detached from the spheroid and invaded the environment (Figure 2a, b).

The MSD in 0d_C_ cells was 0.016 ± 0.004 μm^2^ compared to 4d_C_ cells with 0.012 ± 0.002 μm^2^ (*p* = .012) and 6d_C_ cells with 0.012 ± 0.003 μm^2^ (*p* = .010). The MSD of 4d_C_ and 6d_C_ were similar (*p* = .946) (Figure 3a).

The α was lower of in 0d_C_ cells (0.311 ± 0.298) than in 4d_C_ (0.815 ± 0.243, *p* < .001) and 6d_C_ cells (0.664 ± 0.159, *p* = .003), with a decrease in α in cells from 4d_C_ to 6d_C_, although non-significant (*p* = .173) (Figure 3b). The mitochondrial motion was strongly sub-diffusive in 0d_C_ cells, closely diffusive in 4d_C_ cells, and weakly sub-diffusive in 6d_C_ cells.

In the 4 mg/ml collagen environment, the MSD of the 0d_C_ cells of 0.014 ± 0.005 μm^2^ was slightly larger than for 4d_C_ (0.012 ± 0.004 μm^2^) and 6d_C_ cells (0.011 ± 0.004 μm^2^). These differences were, however, not significant (*p* = .436) (Figure 3a).

The observations for α in 4 mg/ml collagen were similar to those in the 2 mg/ml collagen, although differences were not statistically significant (*p* = .151). The 0d_C_ cells exhibited lower α (0.487 ± 0.284) than the 4d_C_ (0.734 ± 0.220) and 6d_C_ cells (0.659 ± 0.349) (Figure 3b). These α values indicated weaker sub-diffusive motion for the migrated 4d_C_ and 6d_C_ cells than the 0d_C_ cells at the spheroid surface.

### 3.2. Matrix rigidity affects cell stiffness during detachment but not during subsequent migration

The effect of collagen concentration of the matrix on mitochondrial fluctuations and cell stiffness was assessed for varying migration distances from the tumour spheroids.

The increased collagen concentration and rigidity of the matrix did not affect the mitochondrial fluctuations of the cells. The MSD of 0d_C_, 4d_C_ and 6d_C_ cells in 4 mg/ml was similar to that in 2 mg/ml collagen (*p* = .290, *p* = .773 and *p* = .426, respectively) (Figure 3a). An increase, although not significant, in intracellular power-law coefficient α with increased collagen concentration was observed for the 0d_C_ cells at the spheroid surface (*p* = .182) but not for the migrated 4d_C_ (*p* = .444) and 6d_C_ cells (*p* = .964) (Figure 3b).

## 4. Discussion

In the present study, metastatic breast adenocarcinoma cells, MDA-MB-231, were embedded as tumour spheroids in collagen gels of two concentrations (2 and 4 mg/ml collagen) to investigate the mechanical properties of the cancer cells associated with tumour-cell detachment and migration in physiologically relevant 3D environments. Mitochondrial-based particle tracking microrheology was utilised to probe intracellular fluctuations. Cells were distinguished between cells attached to the spheroid, i.e. before tumour-cell detachment, and migrated cells that have employed the necessary processes to detach and gain enhanced migratory properties for cell dissemination away from the tumour spheroid. Cells were denoted according to the distance migrated from the spheroid, normalised to cell diameter (d_C_), where 0d_C_, 4d_C_ and 6d_C_ represented no, medium and high migration, respectively.

This study extends work on the microrheological characterisation of breast cancer cells in 3D environments. MDA-MB-231 cells were used since their mechanics has been extensively studied as single cells in 2D and 3D environments [3, 4] and with and without drug treatment [10, 21]. However, the intracellular mechanics that aid cells in detaching from the tumour and invading the surrounding matrix is not well studied. Here, the microrheology of cells during physiological relevant processes such as tumour detachment and directed migration were characterised in 3D environments.

This study indicates that cells are more solid-like before tumour detachment than after detachment and migration into the surrounding matrix. The cells at the spheroid surface (i.e. 0d_C_) displayed a lower power-law coefficient α, alluding to increased cell stiffness, than migrated 4d_C_ and 6d_C_ cells in both 2 and 4 mg/ml collagen concentrations. In the context of migration, studies have shown an inverse relationship between migratory potential and cell stiffness, irrespective of whether different grades of cancers were from the same cancer type or identical tumour specimens were used [23, 24].

Wullkopf, West (10) reported the stiffness of highly invasive breast cancer (4T1) and pancreatic cancer (KP^R172^HC) cells in the spheroid centre, branch stalk and branch tip during the invasion in compliant and rigid collagen matrices. Here, the spheroid centre, branch stalk and branch tip represented no, medium and large migration, respectively. Irrespective of the matrix rigidity, tip cells that had migrated furthest exhibited lower stiffness than cells in the spheroid centre [10]. These results support the findings in the current study. However, the stiffness of 6d_C_ cells was higher than that of 4d_C_ cells that had migrated a shorter distance. In addition, the cells’ diffusivity α increased from 0d_C_ to 4d_C_ and decreased from 4d_C_ to 6d_C_ irrespective of collagen concentration.

The 0d_C_ cells were attached to the spheroid through cell-cell adhesion, whereas 4d_C_ and 6d_C_ cells detached from the spheroid and migrated away. Hence migrated cells are less mechanically affected by the spheroid and adapt to the matrix rigidity. Interestingly, the cell stiffness at 6d_C_ was higher than at 4d_C_ distance from the spheroid. MDA-MB-231 cells require actomyosin-based contractility to invade a 3D environment despite their spherical morphology [25]. Therefore, it is proposed that the farther migration of 6d_C_ cells compared to 4d_C_ cells requires increased actomyosin contractility and, hence, increased cell stiffness. Furthermore, the mitochondrial fluctuations in migrated 4d_C_ and 6d_C_ cells neared or entered the diffusive regime much earlier than in non-migrated 0d_C_ cells, indicating heightened motor activity to contract the cytoskeleton in preparation for migration and invasion [26].

In the lower collagen concentration (2 mg/ml), the mitochondrial motion was strongly sub-diffusive (α < 0.3) in 0d_C_ cells, diffusive (α = 1.0) in 4d_C_ cells, and weakly sub-diffusive (0.7 < α < 1.0) in 6d_C_ cells. In comparison, the mitochondrial motion was sub-diffusive in 0d_C_ cells and weakly sub-diffusive in 4d_C_ and 6d_C_ cells in the higher collagen concentration (4 mg/ml). These observations indicate that weakly sub-diffusive to diffusive behaviour is characteristic of cells that have migrated away from their spheroid.

The migrating cells exhibited amoeboid motility and moved in a straight path away from the spheroid. Since amoeboid migration is driven by increased myosin II-mediated contractility [25], it is suggested that this contractility mechanism underlies the diffusive mitochondrial motion in migrating cells in this study.

Increased matrix rigidity played a role in decreasing the stiffness of cells at the spheroid surface but did not affect the stiffness of migrated cells. The intracellular power-law coefficient α of 0d_C_ cells was lower in 2 mg/ml than in 4 mg/ml collagen, whereas the α of both 4d_C_ and 6d_C_ cells was similar in 2 and 4 mg/ml collagen matrices. Increased matrix rigidity has been reported to increase intracellular activity and decrease the stiffness of isolated cells in 3D environments [4, 27]. The stiffness of 0d_C_ cells at the spheroid surface is consistent with these reports. For migrated cells, the similar stiffness in soft and stiff matrices for 4d_C_ and 6d_C_ cells, respectively, is consistent with Wullkopf, West (10). They found that the stiffness of detached and migrated invasive breast cancer 4T1 and pancreatic cancer KP^R172^HC cells were similar in collagen matrices of high and low rigidity. In conjunction with the effect of migrated distance, this finding suggests that cells minimise their stiffness during migration irrespective of the matrix rigidity.

The results of this study are consistent with several reports that suggest that decreased cell stiffness mediates the escape of breast cancer cells from the initial metastatic niche [14, 15]. However, this study should be repeated with different cancers to validate the current findings and provide a more comprehensive understanding of the adjustment of intracellular mechanics during cell detachment and invasion.

A target for further research is the interplay of cellular stiffness, ECM remodelling, and biochemical signalling to promote cancer invasion. Matrix metalloprotease (MMP) expression often increases with cancer progression [28], and MMPs affect the remodelling of the ECM by cancer cells, particularly in dense matrices [29]. Transforming growth factor β (TGFβ) promotes cancer invasion at the advanced disease stage [30], e.g. through upregulation of MMPs in breast cancer [8, 31]. Combining cellular microrheology and assessment of MMP and TGFβ expression in spheroid invasion assays will provide more insight into how cancer cells regulate mechanical and biochemical cues to metastasise.

Cell volume is a modulator of cell adhesion and detachment [32]. Wang, Yang (33) found a decrease in volume and motility of metastatic breast cancer cells in a 3D ECM with increasing stiffness but constant adhesion site density. In the current study, the change in collagen concentration affected the pore size and adhesion site density of the collagen fibre network. It is thus of interest to assess the relationship of pore size of the collagen fibre network and cell volume to complement the findings of intracellular mechanics.

## 5. Conclusion

This exploratory study demonstrated that mitochondrial particle tracking microrheology could detect stiffness changes in metastatic breast cancer cells as they detach and migrate away from their spheroids in soft and stiff collagen matrices. The findings revealed the collective effects of enhanced migratory conditions and increased matrix rigidity on the mechanical properties of the cells. The study identified targets for further mechanobiological investigations, including matrix metalloproteases, TGFβ and cell volume, towards therapeutics that target cancer cells at the initial tumour site to inhibit metastatic spread.

## Funding

This research was supported financially by the National Research Foundation of South Africa (grant number IFR14011761118 to TF), the South African Medical Research Council (grant number SIR328148 to TF), the National Institutes of Health [grant numbers U01CA202123 and P01HL120839 to MHZ, Training Grant T32 EB006359 at Boston University that supported JEK), the University of Cape Town (Max & Lillie Sonnenberg International Travel Scholarship and FHS FRC Postgraduate Publication Incentive Award to GH). The funders had no role in study design, data collection and analysis, decision to publish, or preparation of the manuscript. Any opinion, findings, conclusions, or recommendations expressed in this publication are those of the authors and do not necessarily represent the official views of the funding agencies.

## Conflicts of interest statement

The authors declare that they do not have conflicts of interest.

## Supporting information

The data supporting this article can be accessed on the University of Cape Town’s institutional data repository (ZivaHub) under the following doi: http://doi.org/10.25375/uct.13359398.

## Notes

http://doi.org/10.25375/uct.13359398

### Competing Interest Statement

The authors have declared no competing interest.

### Summary of Updates

Reformatted and minor changes

